# FUCCI-based live imaging platform reveals cell cycle dynamics and identifies pro-proliferative compounds in human iPSC-derived cardiomyocytes

**DOI:** 10.1101/2021.12.20.473521

**Authors:** Francesca Murganti, Wouter Derks, Marion Baniol, Irina Simonova, Katrin Neumann, Shahryar Khattak, Kaomei Guan, Olaf Bergmann

## Abstract

One of the major goals in cardiac regeneration research is to replace lost ventricular tissue with new cardiomyocytes. However, cardiomyocyte proliferation drops to low levels in neonatal hearts and is no longer efficient in compensating for the loss of functional myocardium in heart disease. We generated a human induced pluripotent stem cell (iPSC)-derived cardiomyocyte-specific cell cycle indicator system (TNNT2-FUCCI) to characterize regular and aberrant cardiomyocyte cycle dynamics. We visualized cell cycle progression in TNNT2-FUCCI and found G2 cycle arrest in endoreplicating cardiomyocytes. Moreover, we devised a live-cell compound screening platform to identify pro-proliferative drug candidates. We found that the alpha-adrenergic receptor agonist clonidine induced cardiomyocyte proliferation *in vitro* and increased cardiomyocyte cell cycle entry in neonatal mice. In conclusion, the TNNT2-FUCCI system is a valuable tool to characterize cardiomyocyte cell cycle dynamics and identify pro-proliferative candidates with regenerative potential in the mammalian heart.

## Introduction

Heart regeneration in mammals is restricted to the neonatal period when cardiomyocytes can still proliferate (Alkass et al. 2015, Porrello et al. 2011, Ye et al. 2018). Cardiomyocyte proliferation gradually declines after birth and remains at a low level in adulthood (Bergmann et al. 2015, Senyo et al. 2013). Although multiple factors, including changes in metabolism (Honkoop et al. 2019), extracellular matrix (Bassat et al. 2017), and endocrine signaling (Hirose et al. 2019), contribute to this loss of proliferative activity, the exact underlying mechanism that drives cardiomyocyte cell cycle exit still needs further elucidation.

While cardiomyocyte proliferation declines, aberrant cell cycle activity becomes more dominant, leading to multinucleation and endoreplication during the first postnatal weeks (Alkass, Panula, Westman, Wu, Guerquin-Kern and Bergmann 2015). The increase in polyploidy is implicated in the loss of regenerative capacity in postnatal hearts (Hirose, Payumo, Cutie, Hoang, Zhang, Guyot, Lunn, Bigley, Yu, Wang, Smith, Gillett, Muroy, Schmid, Wilson, Field, Reeder, Maden, Yartsev, Wolfgang, Grutzner, Scanlan, Szweda, Buffenstein, Hu, Flamant, Olgin and Huang 2019, Patterson et al. 2017). To date, it is not clear which cues determine the fate of cycling cells and how one can direct nonproductive cell cycle activity to cytokinesis and proliferation. Moreover, this nonproductive cell cycling has made the identification of cardiomyocyte proliferation challenging and has previously led to misinterpretation of the cardiomyocyte renewal capacity (Leone et al. 2015).

The Fluorescence Ubiquitin Cell Cycle Indicator (FUCCI) system is a genetically encoded, protein-based two-color cell cycle indicator that allows for the identification of cycling and noncycling cells and relies on the ubiquitination and degradation of the cell cycle regulators Cdt1 and Geminin (Sakaue-Sawano et al. 2008, Zielke and Edgar 2015). The combination of FUCCI cell cycle indicators with time-lapse microscopy can unequivocally determine whether the outcome of the cardiomyocyte cell cycle is productive (cytokinesis) or nonproductive (endoreplication) (Alvarez et al. 2019, Baniol et al. 2021, Hashimoto et al. 2014, Hashimoto et al. 2015).

Human induced pluripotent stem cell-derived cardiomyocytes (hiPSC-CMs) can currently be robustly generated using established protocols (Burridge et al. 2014), thereby providing an unlimited source of human cardiomyocytes, diminishing the need for cardiomyocytes isolated from rodents, which do not fully recapitulate human physiology. Depending on their level of maturity, hiPSC-CMs still have the capacity to proliferate (Karbassi et al. 2020). Thus, hiPSC-CMs represent an ideal model system to study the kinetics and regulation of human cardiomyocyte proliferation.

Here, we generated a troponin T2-FUCCI (TNNT2-FUCCI) hiPSC line and showed aberrant cell cycle kinetics with G2 arrest in endoreplicating cardiomyocytes compared to those undergoing proliferation and multinucleation. Moreover, we investigated our TNNT2-FUCCI hiPSC-CMs with an autophagy compound library in a live-cell screening approach and identified the alpha-adrenergic agonist clonidine to promote cardiomyocyte proliferation in hiPSC-CMs and cell cycle entry in neonatal mouse cardiomyocytes.

## Methods

### Generation of TNNT2-FUCCI hiPSCs

The pCAG-Fucci2a plasmid (the RIKEN Center for Life Science Technologies, RDB13080) was obtained through RIKEN (Mort et al. 2014, Sakaue-Sawano et al. 2011). mCherry was kindly provided by Roger Y Tsien (Shaner et al. 2004). The Fucci2a portion (mCherry-hCdt1-T2A-mVenus-hGem) was PCR amplified and cloned in frame downstream of the TNNT2 start codon into a custom synthesized (Thermo Fisher Scientific) plasmid backbone containing the ColE1 origin of replication, ampicillin resistance, 1000 bp 5’ and 980 bp 3’ homology arms to the hTNNT2 (NM_000364.4) transcription start site, IRES-Puro and an FRT-flanked PGK-hygromycin selection cassette to generate the targeting vector using the NEBuilder High-Fidelity DNA Assembly Cloning Kit (NEB, E5520). Twenty-one bp downstream of the start codon, including the sgRNA target sites, was excluded from the targeting vector to protect it from CRISPR/Cas9-induced double-strand breaks. The resulting targeting vector was verified by Sanger sequencing, linearized by restriction with SspI and SfiI enzymes and phenol– chloroform-extracted. Two different sgRNAs (*spacer sequences below*), each targeting the region behind the start codon of hTNNT2 that was excluded from the targeting vector, were transcribed *in vitro* using an EnGen® sgRNA Synthesis Kit (NEB, E3322S). The generation and characterization of CRTD5 hiPSCs (hPSCreg, CRTDi005-B) was previously described (Kutsche et al. 2018). CRTD5 hiPSCs (800,000) were electroporated with 10 *μ*g linearized targeting vector, 1 *μ*g of each sgRNA and 60 pmol Cas9-NLS protein (EnGen® Spy Cas9 NLS, NEB, M0646 M) with the Lonza 4D X-unit, pulse CB-150 and Primary Cell 4D-Nucleofector Kit L (V4XP-3024, Lonza). Transfected cells were seeded at clonal density in dishes coated with hESC-qualified Matrigel (Corning Life Sciences, 354277) in mTeSR1 medium (StemCell Technologies, 85850) supplemented for 3 days with 10 *μ*M Rock inhibitor Y-27632 (StemCell Technologies, 78003).

Cells were selected with 50 *μ*g/ml hygromycin B (Thermo Fisher Scientific, 10687010) starting on Day 3 after nucleofection for 7 days, and resistant colonies were manually selected, clonally expanded and screened for correctly targeted clones by colony PCRs amplifying the 5’ and 3’ junction of the targeted alleles from outside of the homology arms into the insert as well as the presence of an intact second allele. Heterozygously targeted clones were selected, and the complete insert was analyzed by Sanger sequencing. The selected complete clone CRTD5-TNNT2-FUCCI #19 was karyotyped by Giemsa banding and showed an intact chromosome set 46,XY[cp20], similar to the parental line (Supp. Fig. 1A).

hTNNT2-sgRNA#3: 5’- GACCATGTCTGACATAGAAG-A3’

hTNNT2-sgRNA#4: 5’- GGTGGTGGAAGAGTACGAGG-3’

### hiPSC culture and maintenance

The hiPSC line (CRTD5) generated from human fibroblasts was used in this study as a control and was obtained from the Stem Cell Engineering facility of the Center for Molecular and Cellular Bioengineering (CMCB), TU Dresden. Research with CRTD5 hiPSC was approved by the *Ethikkommission an der Technischen Universität Dresden* (BO-EK-38012020). Cells were propagated using ReLeSR (StemCell Technologies, #05873) and maintained in mTESR1™ (StemCell Technologies, #85850) on Matrigel™-coated plates (Corning Life Sciences, #354234) under standard culturing conditions (37 °C, 5% CO_2_). Cell cultures were routinely checked for mycoplasma using the LookOut® Mycoplasma PCR Detection Kit (Sigma Aldrich, #D9307).

### hiPSC differentiation into cardiomyocytes

Differentiation of CRTD5 and FUCCI-CRTD5 lines was induced by adaptation of a previously described protocol (Burridge, Matsa, Shukla, Lin, Churko, Ebert, Lan, Diecke, Huber, Mordwinkin, Plews, Abilez, Cui, Gold and Wu 2014, Lian et al. 2013). Briefly, undifferentiated hiPSCs were passaged into 12-well plates using Versene (Thermo Fisher Scientific, #15040066). When hiPSC culture reached 90-95% confluency, differentiation was induced using CDM3 medium (Burridge, Matsa, Shukla, Lin, Churko, Ebert, Lan, Diecke, Huber, Mordwinkin, Plews, Abilez, Cui, Gold and Wu 2014) supplemented with the GSK3β inhibitor CHIR99021 (4 μM, Sigma, #SML1046) for 48 h followed by treatment with IWP2 (5 μM, Tocris, #3533) for an additional 48 h. After, cells were cultured in CDM3 medium. At Day 15 of differentiation, hiPSC-CMs were gently detached from the plate by incubating with 1 mg/ml collagenase B dissolved in CDM3 medium for 30 min at 37 °C. hiPSC-CMs were further dissociated using 0.25% trypsin-EDTA for 5 min at 37 °C. The reaction was stopped by adding a double volume of stop medium (80% CDM3, 20% FBS, ROCK inhibitor). Cells were plated at a density of 1,000,000 cells/well into Matrigel-coated 12-well plates. CDM3 was exchanged every second day until further analysis.

### Imagestream-X and FACS analysis

hiPSC-CMs were detached from the plate by incubating with 1 ml of TrypLE (Thermo Fisher Scientific, #12604013) for 5 min at 37 °C. A double volume of PBS was added to stop the reaction, and the cells were passed through a 100 μm cell strainer. hiPSC-CMs were centrifuged at 200 g for 5 min, fixed with 1% PFA for 20 min and washed 3x with PBS. hiPSC-CMs were incubated with primary antibodies against mVenus (Biorbyt, orb334993, 1:800) and mCherry (Abcam, ab125096, 1:250) and anti-cardiac troponin-T-APC (Miltenyi Biotec, 130-120-403, 1:50) in blocking buffer (PBS, 4% donkey serum, 0.1% Triton X-100 in PBS, 2 mM EDTA) for 2 hours at RT. After washing, the cells were incubated with the secondary antibodies anti-goat Alexa Fluor® 488 (Jackson ImmunoResearch, 705-546-147, 1:500), anti-rabbit Alexa Fluor® 555 (Abcam, ab150062, 1:500) and Hoechst 33342 (Thermo Fisher Scientific, H21492) for 1 h at 4 °C. Cells were then washed 3x with PBS and centrifuged at 200 g for 5 min. Finally, 5 million cells were resuspended in 500 μl of FACS buffer (PBS, 2% FBS, EDTA) and kept on ice until further analysis.

### hiPSC immunohistochemistry

Cells were fixed with 4% formaldehyde solution in PBS for 10 min and stained overnight at 4 °C with primary antibodies against mVenus (1:800, Biorbyt, orb334993) and mCherry (1:250, Abcam, ab125096) and anti-cardiac troponin-T (1:250, Thermo Fisher Scientific, MA5-12960) in blocking buffer (PBS, 4% donkey serum, 0.1% Triton X-100 in PBS, 2 mM EDTA). Cells were then washed 3x with PBS and incubated with the following secondary antibodies in PBS: anti-goat Alexa Fluor® 488 (1:500, Jackson ImmunoResearch, #705-546-147), anti-rabbit Alexa Fluor® 555 (1:500, Abcam, ab150062), anti-mouse Alexa Fluor® 647 (1:500, Jackson ImmunoResearch, #715-606-151) and Hoechst 33342 (Thermo Fisher Scientific, #H3570).

### Measurement of cell area and sarcomere spacing

CRTD5 and FUCCI-CRTD5 cardiomyocytes at Day 25 of differentiation were first costained with antibodies against TNNT2 (mouse, Thermo Fisher Scientific, #MA5-12960) and α-actinin (rabbit, ThermoFisher Scientific, #701914). Imaging of single cardiomyocytes was performed using a Zeiss LSM 700. For cell area analysis, using ImageJ-Fiji software, a defined region of interest (ROI) was defined to outline the outer edges of the cell, and the cell area was measured for each cardiomyocyte.

For sarcomere spacing measurements, an ROI with at least 10 sarcomeres was defined. The intensity or the ROI shows a series of peaks that correspond to the spatial frequency of the sarcomeric pattern, and the amplitude between the peaks was determined to assess the sarcomere spacing.

### Primary cardiomyocyte culture

Whole litters of C57BL/6JRj mice (P0) were used for isolation of neonatal cardiomyocytes. C57BL/6JRj mice were originally obtained from Janvier labs and bred internally in CRTD animal facilities. All procedures were approved by the local ethics committee, Landesdirektion Sachsen (TVT-1/2017). P0 cardiomyocytes were isolated using the Neonatal Heart Dissociation kit (Miltenyi Biotec, 130-098-373) following the manufacturer’s instructions. Cells were seeded in plating medium (20% M199 (Thermo Fisher Scientific, # 12340030), 65% DMEM (Thermo Fisher Scientific, # 31966021), 5% FBS (fetal bovine serum), and 10% HS (horse serum)) at a seeding density of 35,000 cells/well in a 96-well plate coated with Matrigel (Corning Life Sciences, #354234). Cells recovered for 1 day in an incubator (37 °C, 5% CO_2_).

### Live imaging and analysis of TNNT2-FUCCI hiPSCs

HiPSC-derived cardiomyocytes at different time points post cardiomyocyte induction (0, 6 and 30 days) were imaged using a Keyence BZ-X800E microscope (Keyence, Japan). Images were acquired using brightfield, YFP and Cy3 filter sets. For time-lapse imaging, FUCCI hiPSC-derived cardiomyocytes at Day 30 of differentiation were seeded in CDM3 medium at a density of 150,000 cells per well in a 24-well plate (Cellvis, #P24-0-N) coated with Matrigel (Corning Life Sciences, #354234). Time-lapse imaging was performed as previously described (Baniol et al., 2021). Briefly, FUCCI hiPSC-derived cardiomyocytes were imaged every 20 min for 72 h, and the quantification of the mCherry and mVenus fluorescent intensities was performed using Keyence image measurement and analysis software (Keyence, Japan) and ImageJ-Fiji software. A region around the cell was first defined as the background region, and the cell nucleus was segmented using the brightfield signal. The mCherry and mVenus intensities were detected in both the background region and the nucleus. The background signal was subtracted from the nuclear intensity. Single-cell intensity data were aligned based on the peak mVenus intensity and plotted over a period of 40 h using GraphPad Prism.

### Ploidy analysis of cardiomyocytes

Cardiomyocytes were fixed with 4% formaldehyde solution in PBS for 10 min and stained overnight at 4 °C with primary antibodies against TNNI3 (1:500, Abcam, ab56357) and Ki67 (CellSignaling, 9449T) in blocking buffer (PBS, 4% donkey serum, 0.1% Triton X-100 in PBS, 2 mM EDTA). Cells were then washed 3x with PBS and stained with the secondary antibodies anti-goat Alexa Fluor® 488 (Jackson ImmunoResearch, #705-546-147, 1:500), anti-rabbit Alexa Fluor® 555 (1:500, Abcam, ab150062), and Vybrant DyeCycle Violet Stain (Invitrogen, V35003) in PBS. Images were acquired using a Keyence BZ X800 fluorescence microscope (Keyence, Japan) equipped with an imaging cytometer (BZ-H4XI). Image analysis was performed in the open source software CellProfiler 4.2.1 (Carpenter et al. 2006). Nuclei segmentation was performed using the identify primary objects module with an adaptive thresholding method to account for background variances. TNNI3 and Ki67 intensities were measured in segmented nuclei, and thresholds were determined according to their histograms. DNA staining intensities of noncycling (Ki67-negative) cardiomyocyte nuclei (TNNI3-positive) were measured, and ploidy levels were plotted as histograms, from which ploidy thresholds were determined (Supp. Fig. 3).

### Cell plating and culturing and screen conditions

TNNT2-FUCCI hiPSC-CMs were differentiated as described above. hiPSC-CMs were used 30 days after starting the differentiation. hiPSC-CMs were plated on 96-well glass bottom plates (Cellvis, # P96-1-N) coated with Matrigel (Corning Life Sciences, # 354277) at a seeding density of 4000 cells per well. Cells were seeded in a volume of 50 μl RPMI20 medium (RPMI 1640 with 20% FBS). Outer wells were left unused and filled with PBS to exclude well plate edge effects. The medium was changed the next day for RPMI 1640 (Thermo Fisher Scientific, #32404014) + B27 (Thermo Fisher Scientific, #17504044) + 0.1% FBS (Thermo Fisher Scientific, #10500064). Subsequently, hiPSC-CMs were left for 3 days to reduce the baseline level of proliferation induced by the plating medium before starting the screen.

Four days after seeding the cells, the autophagy library (ENZO, #BML-2837-0100) was added to wells by total medium exchange at an end concentration of 25 μM. Then, the medium was supplemented with nontoxic concentrations of Hoechst 33342 (10 ng/ml, Thermo Fisher Scientific, #H3570) and EdU (5 μM, Thermo Fisher Scientific, #C10340). The compound library and controls were divided over two template plates, from which the compounds in the medium were transferred into triplicate screening plates. All pipetting was performed with automated channel pipets to avoid interwell variation. Medium with compounds was not exchanged throughout the 72 h duration of the screen.

### Image acquisition

All images were acquired using a Keyence BZ-X800E compact fluorescence microscope equipped with live imaging cytometer (BZ-H4XI) and CO_2_ control (BZ-H4XT) modules. Live images were acquired using the 10x objective and DAPI, YFP and Cy3 filter sets. Binning was set to 3×3, the gain to 6 dB and exposure times were 28 ms for DAPI, 666 ms for YFP and 167 ms for Cy3 channels. The resulting images were 640×480 pixels. Five nonoverlapping sites were imaged per well. LIVE imaging was performed at Day 0 directly after adding the compound library and every 24 h until 72 h.

After the last LIVE acquisition, hiPSC-CMs were fixed by incubation with 4% formaldehyde solution in PBS for 15 min. Subsequently, an EdU click-it reaction (Thermo Fisher Scientific, #C10340) was performed according to the manufacturer’s protocol to visualize EdU incorporation. Images of fixed cells were acquired using DAPI and Cy5 channels, binning was set to 3×3, gain to 6 dB and exposure times were 100 ms for DAPI and 10 ms for Cy3.

### Automated image analysis

Automated image analysis was performed in the open source software CellProfiler 4.2.1 (Carpenter, Jones, Lamprecht, Clarke, Kang, Friman, Guertin, Chang, Lindquist, Moffat, Golland and Sabatini 2006). Briefly, single channel images from DAPI, YFP and Cy3 channels were imported into the program. In the first module, the YFP images were enhanced to reduce uneven backgrounds. Subsequently, the DAPI channel was used to identify all nuclei and assign them as primary objects. In the next modules, the intensity of mVenus and mCherry from the YFP and Cy3 channel images within the nuclei was measured, and a threshold was set to categorize the nuclei as positive (+) or negative (−) for these channels. In the following modules, we assigned all nuclei to a single category as follows: mVenus- and mCherry-nuclei: BLUE; mCherry^+^ and mVenus^−^ nuclei: RED; mCherry^+^ and mVenus^+^ nuclei: YELLOW; mVenus^+^ and mCherry^−^ nuclei: GREEN. In the last module, counts were exported into spreadsheets for further data analysis.

### Data analysis

Data analysis was performed using the open-source data analytics software KNIME (Konstanz Information Miner) version 4.3.1. The general approach for screen analysis was followed as described by Stöter et al. (Stoter et al. 2019). Briefly, CellProfiler output files were imported, and the following steps were performed: quality control, filtering, grouping, normalization and statistics. As a first step, data from individual sites were excluded from analysis if the number of nuclei was far below (>2 STDEV) mean levels (indicating problems with focus, e.g.,). In the second step, the results from imaging sites were grouped by wells, giving mean values per well from the five sites imaged. In the next steps, the data from wells were matched with locations on the plates and the compound information. The percentages per compound were normalized to the control (=100%). Next, wells were grouped by treatment, and if parameters showed STDEVs larger than half the value of the parameter, they were excluded as quality controls. Data were exported into Excel, and the main parameter (% of mVenus^+^ in all FUCCI^+^) was plotted using GraphPad Prism software (Fig. 3B).

### Clonidine treatment in neonatal mouse pups

Animals were housed in the Comparative Medicine Biomedicum (Karolinska Institutet, Stockholm) animal facility on a 12-h light/dark cycle and were provided food and water ad libitum. All breeding and organ collection protocols were performed in accordance with the Swedish and European Union guidelines and approved by the institutional ethics committee (Stockholms Norra Djurförsöksetiska Nämnd). C57BL/6N neonatal mice were injected subcutaneously with a volume of 0.1 ml of PBS with EdU (20 mg/kg, Invitrogen, # E10187) and clonidine (60 ng/pup, Sigma, # C7897). Clonidine was given for an estimated weight of 1.5 g/pup throughout the experiment; therefore, a fixed dose of 60 ng/pup was used. Neonatal mice were sacrificed at P7 by decapitation, and hearts were dissected and collected in PBS, cryoprotected in 30% sucrose and flash-frozen in isopentane.

## Results

### Generation and validation of TNNT2-FUCCI in hiPSC-derived cardiomyocytes

We generated a TNNT2-FUCCI hiPSC line using a CRISPR–Cas9 approach (see Methods). TNNT2-FUCCI hiPSCs expressed the FUCCI construct under the control of the cardiomyocyte-specific TNNT2 promoter (Fig. 1A). The generated TNNT2-FUCCI hiPSC line showed a normal karyotype, and pluripotency characterization showed high expression of bona fide pluripotency markers and could generate three germ layers (Sup. Fig. 1A-C). FUCCI fluorescence (mCherry/mVenus) became visible from Day 6 post differentiation into cardiomyocytes (Fig. 1A) when TNNT2 expression was first detected (Burridge PW et al., 2011). At Day 30, in most cells, an FUCCI signal was detected (Fig. 1A), and all FUCCI-expressing cells were TNNT2 positive (Fig. 1B). To further confirm the cardiomyocyte-specific expression of TNNT2-FUCCI fluorescence, we used imaging flow cytometry (Amnis imagestream-X analyzer). We analyzed 50,000 cells defined by their TNNT2 expression and found that 85.4% were mCherry positive, 5.1% were mVenus positive and 1.6% were double positive cardiomyocytes (Fig. 1C). Importantly, we found no FUCCI expression in the TNNT2-negative cell fraction. However, some signal in the TNNT2-negative fraction detected in the green and red fluorescence channels could be assigned to autofluorescence (Supp. Fig. 2A).

**Figure 1.**
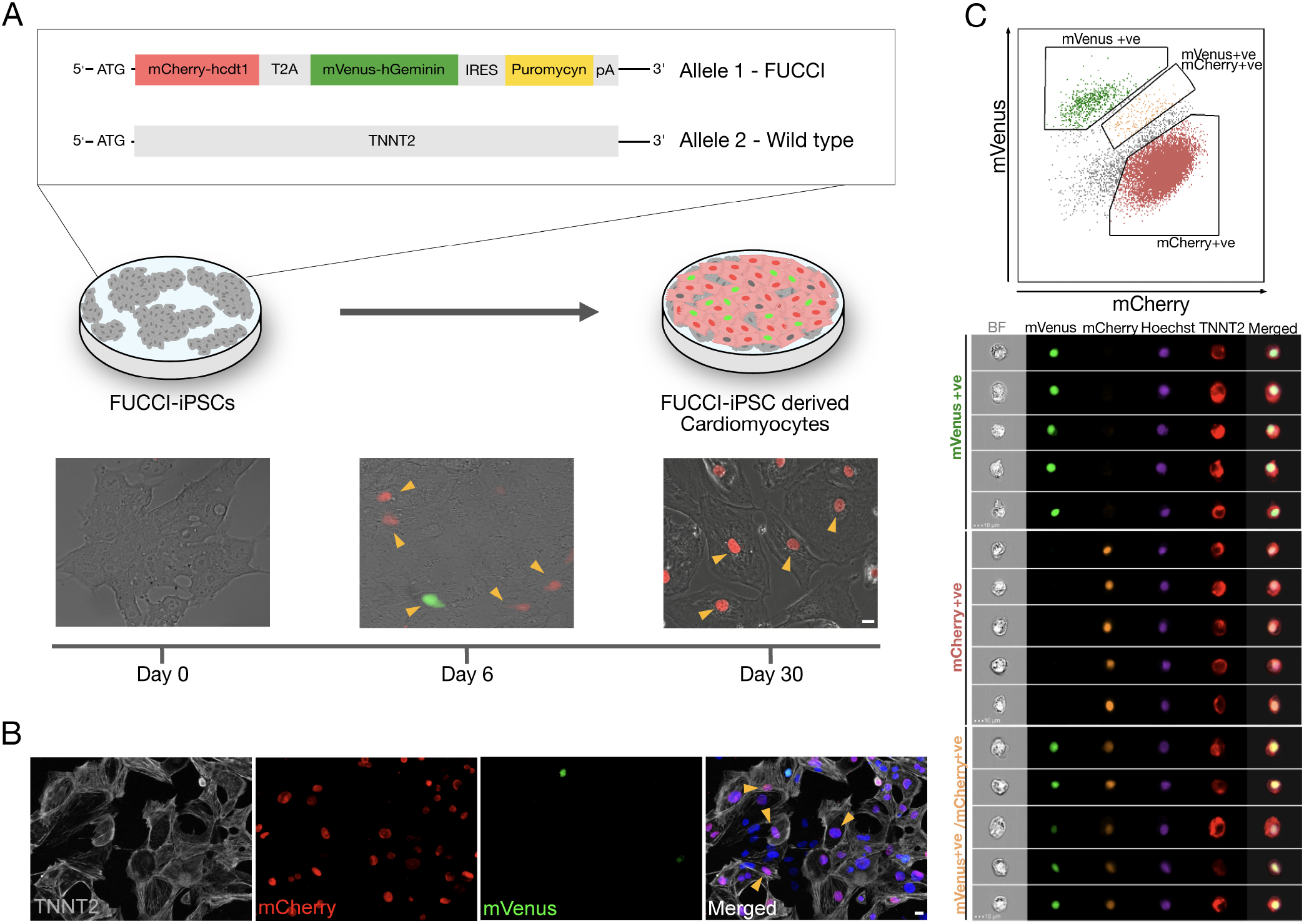
Generation and validation of TNNT2-FUCCI in hiPSC-CMs. **A.** Schematic representation showing TNNT2-FUCCI reporter cell line generation and differentiation toward cardiomyocytes. Upon differentiation into cardiomyocytes, the FUCCI signal became visible at Day 6. Arrowheads indicate FUCCI-positive nuclei. Scale bar, 20 μm. **B.** Immunocytochemistry showing cardiomyocytes at Day 30 of differentiation. Arrowheads indicate cardiomyocytes (TNNT2-positive) that were FUCCI-positive, demonstrating the specificity of the FUCCI reporter. Scale bar, 20 μm. **C.** Imaging flow cytometry shows TNNT2-positive cells expressing nuclear mVenus and/or mCherry, indicating cardiomyocyte specificity of the TNNT2-FUCCI system.

To demonstrate that the TNNT2-FUCCI signal reliably indicates the cell cycle status of cardiomyocytes, we costained FUCCI cardiomyocytes with cyclin-dependent kinase-1 (CDK1) and found a complete overlap of mVenus-positive and CDK1-positive nuclei, supporting that mVenus-positive cardiomyocytes indeed represent cells in the S/G2/M phase of the cell cycle (Supp. Fig. 2B).

TNNT2-FUCCI cardiomyocytes showed a similar sarcomere spacing pattern (1.97 μm ± 0.06 μm SEM) and cell size (2525.4 μm^2^ ± 259.2 μm^2^ SEM) as controls (Supp. Fig. 2C and 2D), suggesting that the integration of the TNNT2-FUCCI knock-in did not compromise the functionality of the TNNT2-FUCCI cardiomyocytes.

In summary, we showed that TNNT2-FUCCI expression is limited to cardiomyocytes and reliably detects cell cycle progression.

### Live cell imaging of TNNT2-FUCCI cardiomyocytes shows differences in cell cycle progression with G2 phase arrest in polyploidy

We tracked single TNNT2-FUCCI cardiomyocytes over several days and plotted the FUCCI oscillation pattern in cardiomyocytes undergoing cell division, multinucleation and polyploidization (Fig. 2A to 2C, and supplementary movies S1 to S3). In all three cycling populations, mVenus fluorescence started to increase concomitant with the reduction of the mCherry signal, marking the end of the G1 phase and the beginning of the S/G2 phase. In early mitosis, mVenus fluorescence dropped sharply concomitant with nuclear envelope breakdown in dividing and multinucleating TNNT2-FUCCI cardiomyocytes (Fig. 2A and 2B). The average duration of the S/G2/M phase was 16.38 h ± 0.84 h SEM in dividing cells and 17.29 h ± 0.55 h SEM in multinucleating cells, with no significant difference between the two populations (p>0.05, Fig. 2D). In contrast, polyploid cells showed a different FUCCI oscillation pattern. An increase in the mCherry signal was detected before the loss of mVenus, which dropped slowly, suggesting G2 phase arrest without activation of the anaphase-promoting complex (Ballabeni et al. 2004). Accordingly, no mitotic features, such as cell rounding or cytoplasmic localization of mVenus, were detected (Fig. 2C). Moreover, mVenus expression was detected over a significantly longer period in polyploid cells compared to dividing (p=0.0009) and multinucleating (p=0.0029) cells, with an average duration of the S/G2 phase of 24.5 h ± 2.77 h SEM (Fig. 2D).

**Figure 2.**
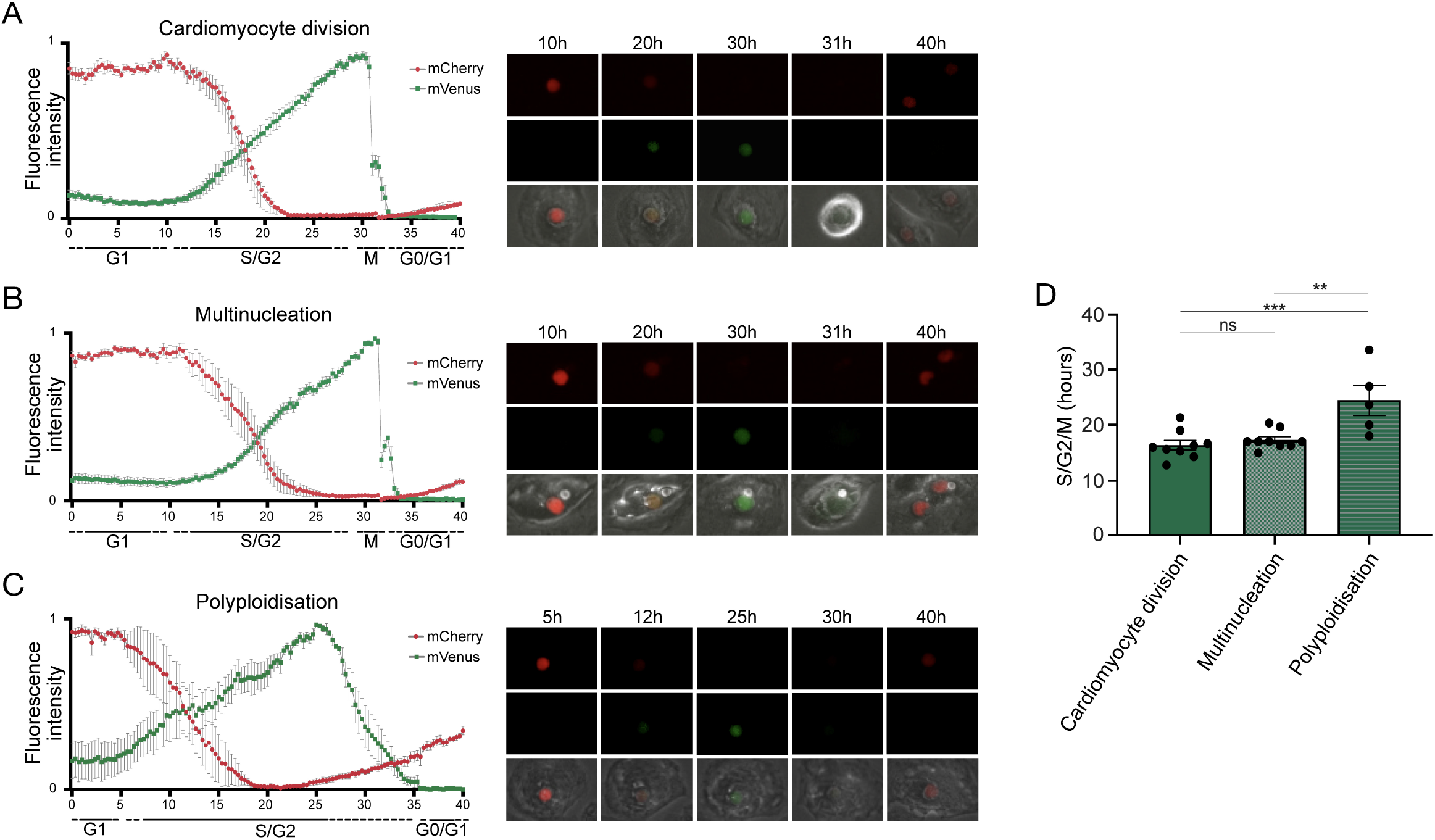
Quantification of cell cycle duration using live cell imaging of hiPSC-CMs. **A-C** Oscillation patterns of FUCCI fluorescence measured by live imaging in cardiomyocytes undergoing (A) cell division (n=9), (B) multinucleation (n=9) and (C) polyploidization (n=5). Data show mean intensity ± SEM. **D.** Quantification of the total duration of the combined S, G2 and M phases shows a significant increase in cell cycle duration in cardiomyocytes undergoing polyploidization. Data show mean intensity ± SEM. P value was determined by one-way ANOVA. * p<0.05, **p<0.01, ***p<0.001.

These data suggest that hiPSC-derived cardiomyocytes that become polyploid do not enter mitosis and remain arrested in the G2 phase.

### Live image-based TNNT2-FUCCI screening identifies cell cycle activators

Next, we devised a live screening platform in which we probed TNNT2-FUCCI cardiomyocytes for cell cycle entry using a library of 94 autophagy-related compounds (Fig. 3A, and Methods). TNNT2-FUCCI live cardiomyocytes were imaged, and mVenus and mCherry nuclear fluorescence was documented at 24 h, 48 h, and 72 h (Fig. 3). Proproliferative activity was determined as the percentage of mVenus^+^ (S/G2/M phase) cardiomyocyte nuclei and normalized to the control group at all three time points (Fig. 3 and Supp. Fig. 3A and 3B). The acquisition time point of 48 h with the smallest coefficient of variance (24 h, 48 h, 72 h; 13.1%, 12.1%; 28.0%, respectively) was chosen to select six pro-proliferative candidates. We selected the six pro-proliferative candidate compounds based on their increase in the percentage of mVenus^+^ nuclei and their biological significance (see Methods and Supp. Fig. 3).

**Figure 3.**
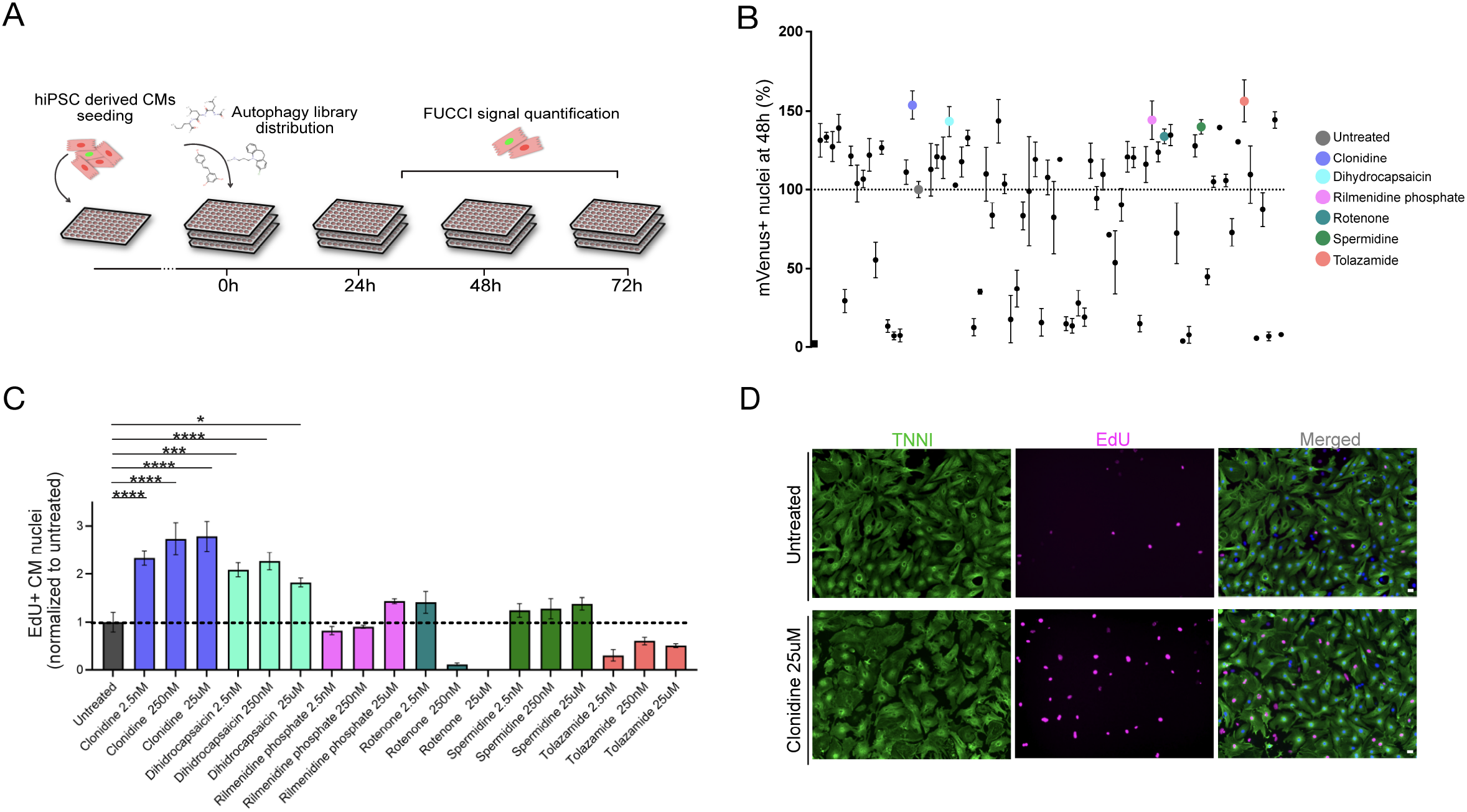
Live-image-based FUCCI hiPSC-CM cell cycle activation screen identifies compounds that induce cell cycle activity. **A.** Schematic drawing of the screening experiment. hiPSC-CMs were seeded in 96-well plates and stimulated with 94 compounds of an autophagy-related compound library at a concentration of 25 μM. FUCCI signal was documented at 24 h, 48 h and 72 h. **B.** The effect of each compound is shown as the percentage of mVenus+ nuclei relative to the control at the 48 h time point. Error bars represent STDEV between triplicates. Compounds found to increase the percentage of mVenus+ myocyte nuclei that were subjected to further validation are indicated in color. **C.** Selected compounds were further tested in mouse neonatal cardiomyocytes (mNCMs) at three different concentrations (2.5 nM, 250 nM, 25 μM). Cell cycle activity was validated by EdU incorporation and subsequent detection at 72 h, n=3 per sample. Values are mean ± SEM. P values were determined by one-way ANOVA. * p<0.05, **p<0.01, ***p<0.001, ****p<0.0001. **D.** Representative immunocytochemistry images of untreated and clonidine-treated mNCMs with EdU incorporation detected in magenta and cardiac troponin T in green. Scale bar, 20 μm.

### Validation of pro-proliferative candidate compounds in neonatal cardiomyocytes

The six selected compounds were tested at three different concentrations to establish potential concentration dependencies (2.5 nM, 250 nM, 25 μM) in mouse neonatal cardiomyocytes (mNCMs). Cell cycle activation was validated by EdU incorporation after 72 h (Fig. 3C). We found two compounds that significantly increased cell cycle activity at all tested concentrations. Of these, the alpha-adrenergic agonist clonidine showed the most pronounced concentration-dependent effect (2.78-fold increase compared to untreated, p<0.0001) on cell cycle activity in mNCMs (Fig. 3C and 3D and Supp. Fig. 3C). Hence, we continued to explore the pro-proliferative potential of clonidine.

### Clonidine triggers proliferation in hiPSC-derived cardiomyocytes and cell cycle activity in mouse neonatal cardiomyocytes

We assessed whether clonidine-induced cell cycle activity results in proliferation or polyploidy in hiPSC-derived cardiomyocytes (Fig. 4A to E). Cell cycle activity measured through Ki67 expression increased from 9.0% ± 1.23% to 22.0% ± 4.51% 72 h after treatment in hiPSC-CMs (p=0.03, Fig. 4A). Clonidine did not cause any changes in polyploidy (p>0.05, Fig. 4B, Supp. Fig. 4A) or in binucleation (p>0.05, Fig. 4C), suggesting that most clonidine-induced cell cycle activity results in proliferation. Consistent with these findings, we found an increase in aurora B kinase (AurKB)-positive midbodies, indicative of late phase cytokinesis, from 0.83% ± 0.17% in untreated hiPSC-derived cardiomyocytes to 1.69% ± 0.24% in clonidine-treated hiPSC-derived cardiomyocytes (p=0.04, Fig. 4D and E).

**Figure 4.**
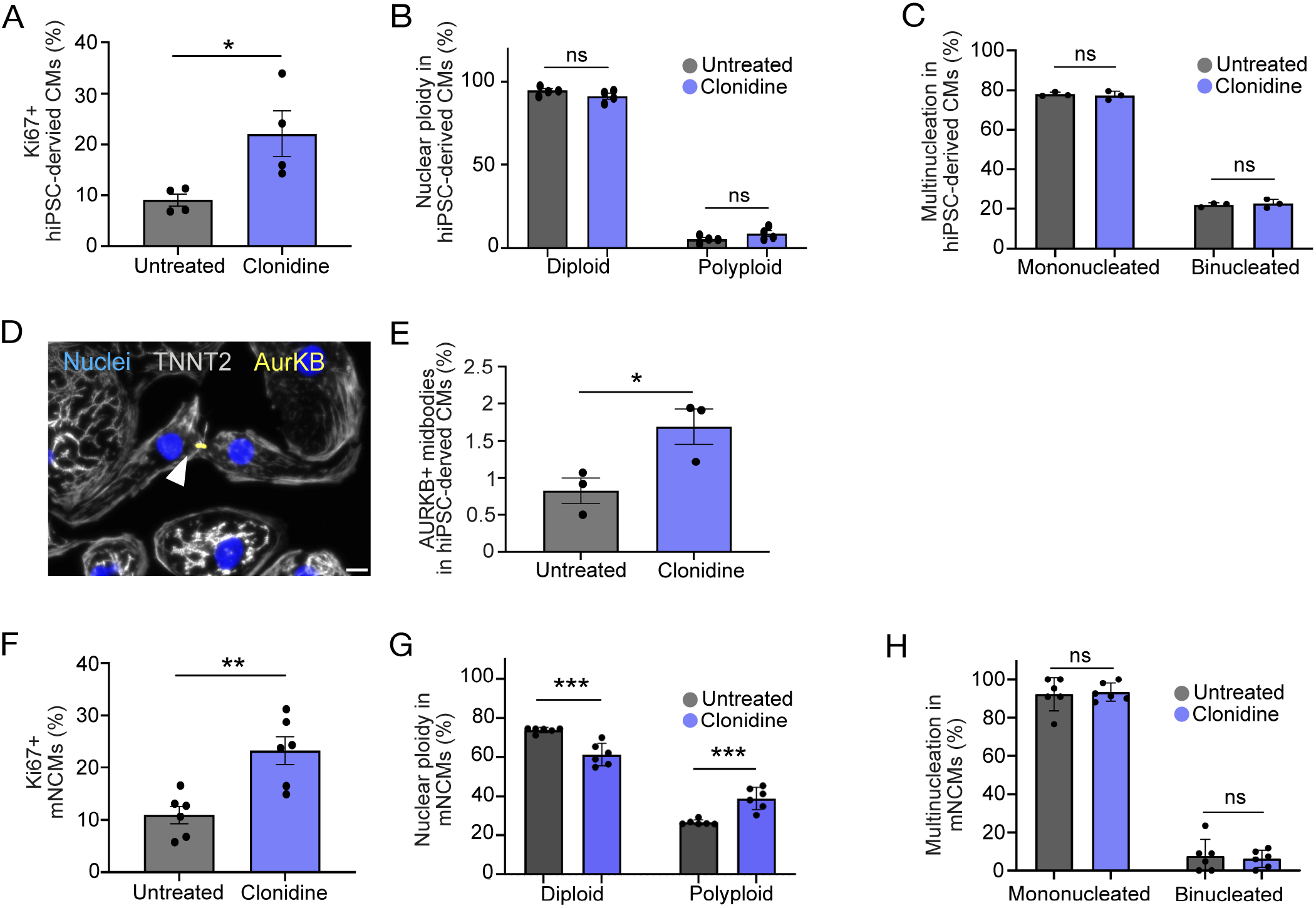
Effect of clonidine on cell cycle activity and proliferation in hiPSC-CMs and mNCMs. **A.** Quantification of Ki67-expressing cardiomyocytes after 72 h of clonidine treatment in hiPSC-CMs. P values were determined by unpaired t test. **B-C.** Cell cycle activity induced by clonidine did not show any changes in (B) nuclear ploidy or (C) binucleation. P values were determined by two-way ANOVA. **D-E.** Aurora B kinase-positive midbodies, indicative of late phase cytokinesis, increased significantly in clonidine-treated hiPSC-CMs. Scale bar, 20 μm. P values were determined by unpaired t test. **F.** Clonidine significantly increased the percentage of Ki67^+^ cardiomyocytes in mNCMs. P values were determined by unpaired t test. **G.** Quantification of nuclear ploidy in mNCMs showed an increase in nuclear ploidy. P values were determined by two-way ANOVA. **H.** Ratios of binucleated mNCMs were not altered by clonidine treatment. P values were determined by two-way ANOVA. Values are mean ± SEM. * p<0.05, **p<0.01, ***p<0.001.

Next, we investigated whether clonidine elicits similar effects in mNCMs that are more restricted in their capacity to proliferate. We found an increase in cell cycle activity, measured through Ki67 expression, from 10.97% ± 1.68% SEM in untreated mNCMs to 23.23% ± 2.63% SEM in clonidine-treated mNCMs (p=0.002, Fig. 4F). The ratio of binucleated cardiomyocytes was not altered by clonidine treatment (p>0.05, Fig. 4H), but we found an increase in nuclear ploidy with an increase in the tetraploid fraction from 26.52% ± 0.54% SEM in untreated mNCMs to 38.93% ± 2.32% SEM in clonidine-treated mNCMs (p=0.0004, Fig. 4G, Supp. Fig. 4B), suggesting that a substantial proportion of clonidine-induced cell cycle activity can be attributed to nuclear polyploidy. Additionally, we performed AurKB staining on clonidine-treated and control mNCMs. Although we observed AurKB-positive cardiomyocyte nuclei in all cultures (Supp. Fig. 5), we could not detect any positive midbodies in these cultures (more than 10,000 cardiomyocytes analyzed), suggesting that clonidine does not stimulate cytokinesis in cell cycle-active mNCMs.

### Clonidine induces cell cycle activity in the neonatal mouse heart

To explore whether clonidine triggers cell cycle activity in the mouse heart, similar to what we found *in vitro*, we administered clonidine to neonatal mice. Clonidine was given along with EdU daily from P1 to P5 (60 ng/day), and the hearts were collected for analysis at P7 (Fig. 5A). The number of EdU^+^ cardiomyocyte nuclei (EdU^+^/PCM-1^+^) was significantly increased in the clonidine-treated group (36.15% ± 2.72%) compared to control animals (25.54% ± 1.80%) (p=0.02, Fig. 5B and 5C), demonstrating that clonidine triggers cardiomyocyte cell cycle activity in both *in vitro* and *in vivo* mouse neonatal hearts.

**Figure 5.**
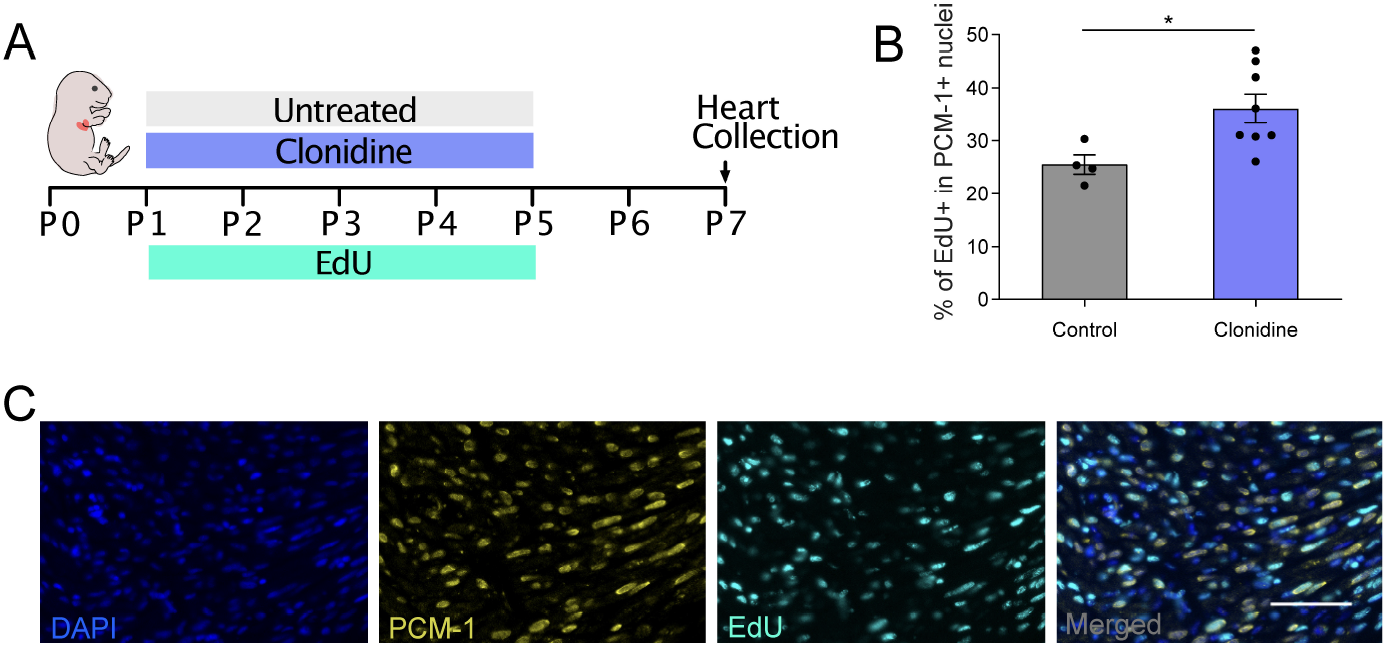
Clonidine induces cell cycle activity in neonatal mice. **A.** Neonatal mice were subjected to 4 days (P1-P5) of clonidine (60 ng/pup) and EdU treatment (20 mg/kg); pups were sacrificed, and hearts were collected at P7. **B-C.** Quantification of EdU^+^ cardiomyocyte (PCM-1^+^) nuclei in sectioned mouse hearts showed an increase in EdU incorporation in clonidine-treated neonatal mice. Values are mean ± SEM. P value was determined by unpaired t test. * p<0.05. Scale bar, 50 μm.

## Discussion

Heart failure is among the leading causes of death in the Western world, and currently available treatment is limited to salvaging existing cardiomyocytes or heart transplantation. Augmenting the proliferation of existing cardiomyocytes is often proposed as a promising future strategy for reverting disease progression (Tzahor and Poss 2017). To achieve this goal, detailed knowledge of the human cardiomyocyte cell cycle and how it can be manipulated is crucial. Here, we generated TNNT2-FUCCI cardiomyocytes to reveal cell cycle kinetics in human cardiomyocytes undergoing proliferation, binucleation and polyploidization. To show the versatility of TNNT2-FUCCI, we devised a live cell screening platform to assess the pro-proliferative effects of compounds from an autophagy library. Using this platform, we identified clonidine as a compound that induces cell cycle activity in cardiomyocytes, resulting in the proliferation of hiPSC-CMs.

### Generation and characterization of a hiPSC line with the cardiomyocyte-specific TNNT2-FUCCI reporter

The identification of cycling cardiomyocytes is critical for studying cardiomyocyte proliferation (Derks and Bergmann 2020, Leone, Magadum and Engel 2015). Here, we generated the TNNT2-FUCCI hiPSC reporter line and differentiated it into cardiomyocytes. Consistent with previous data, we showed that inactivation of one TNNT2 locus by FUCCI knock-in did not cause structural impairments in cardiomyocytes (Ahmad et al. 2008). After 6 days in culture, we found the first cells expressing TNNT2-FUCCI along with the appearance of beating cardiomyocytes at Day 8 of differentiation (Burridge, Matsa, Shukla, Lin, Churko, Ebert, Lan, Diecke, Huber, Mordwinkin, Plews, Abilez, Cui, Gold and Wu 2014). We found a complete overlap of cyclin-dependent kinase 1 (CDK1), which oscillates in the cell cycle and shows activity in S/G2/M phases (Bashir and Pagano 2005), with mVenus-geminin-positive cardiomyocytes, verifying the fidelity of geminin expression and thereby with the TNNT2-FUCCI system.

### Single-cell live imaging of cycling cardiomyocytes

Video time-lapse microscopy is considered the gold standard to unequivocally determine the outcome of the cardiomyocyte cell cycle as division, multinucleation or polyploidization (Derks and Bergmann 2020, Leone and Engel 2019, Leone, Magadum and Engel 2015). Here, we combined time-lapse microscopy with our TNNT2-FUCCI sensor to determine the length and distinct phases of the cardiomyocyte cell cycle. We found that the S/G2/M phase took approximately 18 h for both dividing and binucleating cells with similar TNNT2-FUCCI fluorescent oscillation patterns, documenting a slightly longer cell cycle length than in mouse embryonic E14.5 cardiomyocytes (10.4 hours) (Hashimoto, Yuasa, Tabata, Tohyama, Hayashiji, Hattori, Muraoka, Egashira, Okata, Yae, Seki, Nishiyama, Nakajima, Sakaue-Sawano, Miyawaki and Fukuda 2014) and in mouse neonatal cardiomyocytes at P0 (15.1 hours) (Baniol, Murganti, Smialowska, Panula, Lázár, Brockman, Giatrellis, Derks and Bergmann 2021). In contrast, human cardiomyocytes undergoing polyploidy showed a longer S/G2/M phase duration of approximately 25 h and prominent changes in the TNNT2-FUCCI fluorescence intensity pattern, exemplifying the difficulty in estimating annual cell cycle activity rates based on a fixed duration of cell cycle length (Bergmann and Jovinge 2014, Bergmann et al. 2011). Furthermore, our data suggest that polyploidy is not only a result of karyokinesis failure (Han et al. 2020) because we did not observe nuclear envelope breakdown in cardiomyocytes but can also be a result of G2-phase arrested cycling cardiomyocytes.

### Live TNNT-FUCCI screening platform

Here, we developed a fluorescence-based live imaging screening platform based on the TNNT2-FUCCI system. This screening platform does not require immunofluorescence staining and eliminates the need for total cell counts and the incorporation of nucleotide analogs (e.g., BrdU and EdU). Live imaging of cycling cardiomyocytes allows for multiple imaging time points to establish the efficacy time course of pro-proliferative compounds. Due to the cardiomyocyte specificity of the TNNT2-FUCCI system, we eliminated false-positive detection of proliferative events originating from nondepleted cycling noncardiomyocytes. Moreover, TNNT-FUCCI cardiomyocyte specificity allows sophisticated cocultures of cardiomyocytes with other cell types and could even be combined with recently developed cardiac organoids (Hofbauer et al. 2021). Our platform can be used with any type of library to identify pro-proliferative candidates. In this study, we subjected TNNT2-FUCCI hiPSC-derived cardiomyocytes to a library of compounds regulating autophagy. Autophagy has been implicated in heart regeneration (Xie et al. 2021) and exerts numerous roles in myocardial stress responses (Delbridge et al. 2017). Autophagy reduces cellular stress caused by reactive oxygen species, which in turn plays a role in cardiomyocyte cell cycle exit (Puente et al. 2014).

### Validation of screen-identified compounds in mouse cardiomyocytes

We selected six compounds for further validation in mouse neonatal cardiomyocytes (Supp. Table 1), which are considered more mature than iPSC-derived cardiomyocytes (Karbassi, Fenix, Marchiano, Muraoka, Nakamura, Yang and Murry 2020), with a partial loss of their capacity to proliferate. We found that only two of the six compounds significantly enhanced cell cycle activity, potentially due to higher levels of maturation in mouse NCMs compared to hiPSC-CMs and cross-species variation in cardiomyocyte physiology. One of these two compounds was dihydrocapsaicin, an analog of the active component of chili pepper with documented effects on cardiomyocytes, including autophagy induction (Wei et al. 2020) and attenuation of mitochondrial function (Qiao et al. 2020).

The most robust and dose-dependent effect on cardiomyocyte cell cycle entry, however, was elicited by the alpha-adrenergic agonist clonidine. Alpha-adrenergic receptors are predominantly found on smooth muscle cells of blood vessels, mediating vasoconstriction, but can also be detected on cardiomyocytes (Woodcock et al. 2008). Clonidine also binds and activates the imidazoline receptor, leading to a decrease in the level of cAMP in cells, resulting in downstream autophagy activation (Williams et al. 2008). Although it is unclear whether clonidine elicits its pro-proliferative effects via autophagy induction or other mechanisms, we chose to further investigate whether clonidine treatment in addition to cell cycle entry also results in successful mitosis and cytokinesis. Based on detected AurKB-positive midbodies, clonidine promotes cytokinesis in hiPSC-CMs. While the increase in cell cycle activity did not alter the ratio between diploid and polyploid cells in hiPSC-CMs, in mouse NCMs, we did observe an increase in polyploid CMs and no AurKB-positive midbody formation after clonidine treatment. The latter indicates that clonidine-induced cell cycle activity was not productive and that for successful completion of mitosis and cytokinesis in mouse NCMs, additional stimuli are required.

In conclusion, we generated TNNT2-FUCCI hiPSCs and demonstrated that TNNT2-FUCCI hiPSC-derived cardiomyocytes enable analysis of cell cycle entry and progression, providing a powerful platform for screening and validation of pro-proliferative candidates in human cardiomyocytes.

## Acknowledgments

The CRTD5-TNNT2-FUCCI reporter hiPSC line was generated by the Stem Cell Engineering Facility, a core facility of the Center for Molecular and Cellular Bioengineering (CMCB) at Technische Universität Dresden. Giemsa staining and analysis of metaphase chromosomes were performed at the Molecular Cytogenetics lab at Jena University Hospital. We thank Marc Bickle from MPI-CBG in Dresden for advice on image acquisition strategies and for helping to set up the analysis in KNIME. We thank the Light Microscopy and Flow Cytometry core facilities of BIOTEC/CRTD for their help with imaging and flow cytometry analyses. O.B. was supported by the Center for Regenerative Therapies Dresden, the Karolinska Institute, the Swedish Research Council, the Ragnar Söderberg Foundation, the Åke Wiberg Foundation, and the LeDucq Foundation.

## Author contributions

Conceptualization, F.M., W.D., and O.B.; Methodology, F.M., M.B., W.D., I.S.; Investigation, F.M., M.B., W.D., K.N., S.K., and O.B.; Writing – Original Draft, F.M., W.D. and O.B.; Writing – Review & Editing, F.M., W.D., K.G., and O.B.; Supervision, F.M., W.D., K.G., and O.B.; Funding Acquisition, O.B.

---

**Supplemental Fig. 1. Characterization of TNNT2-FUCCI hiPSC line**

**A.** The CRTD5-TNNT2-FUCCI hiPSC line was karyotyped by Giemsa banding and showed an intact chromosome set similar to the parental line. **B.** ICC shows the capacity to differentiate into the three germ layers; markers are shown in green. **C.** Flow cytometry analysis showing high levels of expression of the pluripotency markers OCT4, SOX2, SSEA4 and TRA1-60. Blue markers indicate no antibody control, and red indicates the antibody-stained sample. The percentages shown refer to antibody-stained sample.

**Supplemental Fig. 2. Characterization of TNNT2-FUCCI cardiomyocytes**

**A.** Imaging flow cytometry recordings of nonmyocytes. All signals observed in the TNNT2-negative fraction in the green and red fluorescence channels could be assigned to autofluorescence. **B.** Costaining of FUCCI cardiomyocytes with cyclin-dependent kinase-1 (CDK1) shows an overlap of mVenus-positive and CDK1-positive nuclei. Scale bar, 20 μm. **C.** TNNT2-FUCCI cardiomyocytes showed similar sarcomere spacing patterns, as determined using TNNT2 and α-actinin staining. Scale bar, 20 μm. **D.** Cell size was not compromised in FUCCI cardiomyocytes. Scale bar, 20 μm.

**Supplemental Fig. 3. Autophagy compound screen of TNNT2-FUCCI cardiomyocytes**

**A-B.** The effect of each compound is shown as the percentage of mVenus+ nuclei relative to the DMSO control (=100%). Timepoints of 24 h and 72 h were used to generate these graphs. Error bars represent STDEV between triplicates. **C.** Quantification and representative immunocytochemistry images of untreated and clonidine (25 μM and 100 μM)-treated mNCMs with EdU incorporation in purple and cardiac troponin T in green. Scale bar, 50 μm. Values are mean ± SEM. * p<0.05, **p<0.01.

**Supplemental Fig. 4. Ploidy of noncycling cardiomyocytes in neonatal mouse cardiomyocytes**

Histograms showing the DNA intensity distribution of noncycling cardiomyocyte nuclei (TNNI3-positive/Ki67-negative) of (A) hiPSC-derived cardiomyocytes and (B) neonatal mouse cardiomyocytes.

**Supplemental Fig. 5. Aurora B staining of neonatal cardiomyocytes**

Nuclear aurora B staining (red) indicates G2/M activity of neonatal cardiomyocytes (green, arrowheads). More than 10,000 cardiomyocytes were analyzed, but no aurora B-positive midbody assembly at the site of abscission was detected.

**Supplementary Table 1. Characterization of candidate compounds.**

