## Supplemental Fig. S1-S5 for "FUCCI-based live imaging platform reveals cell cycle dynamics and identifies pro-proliferative compounds in human iPSC-derived cardiomyocytes"

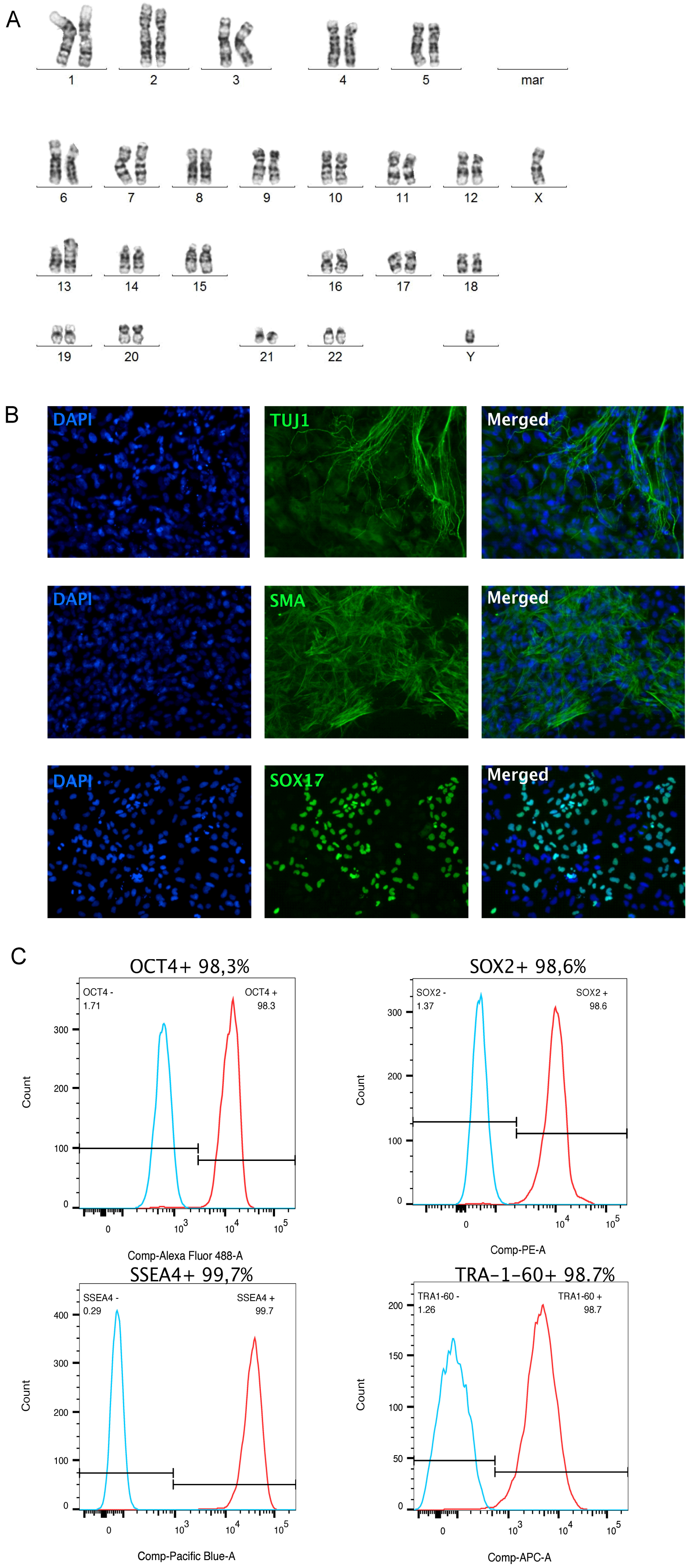


SUPPLEMENTARY FIGURE 1

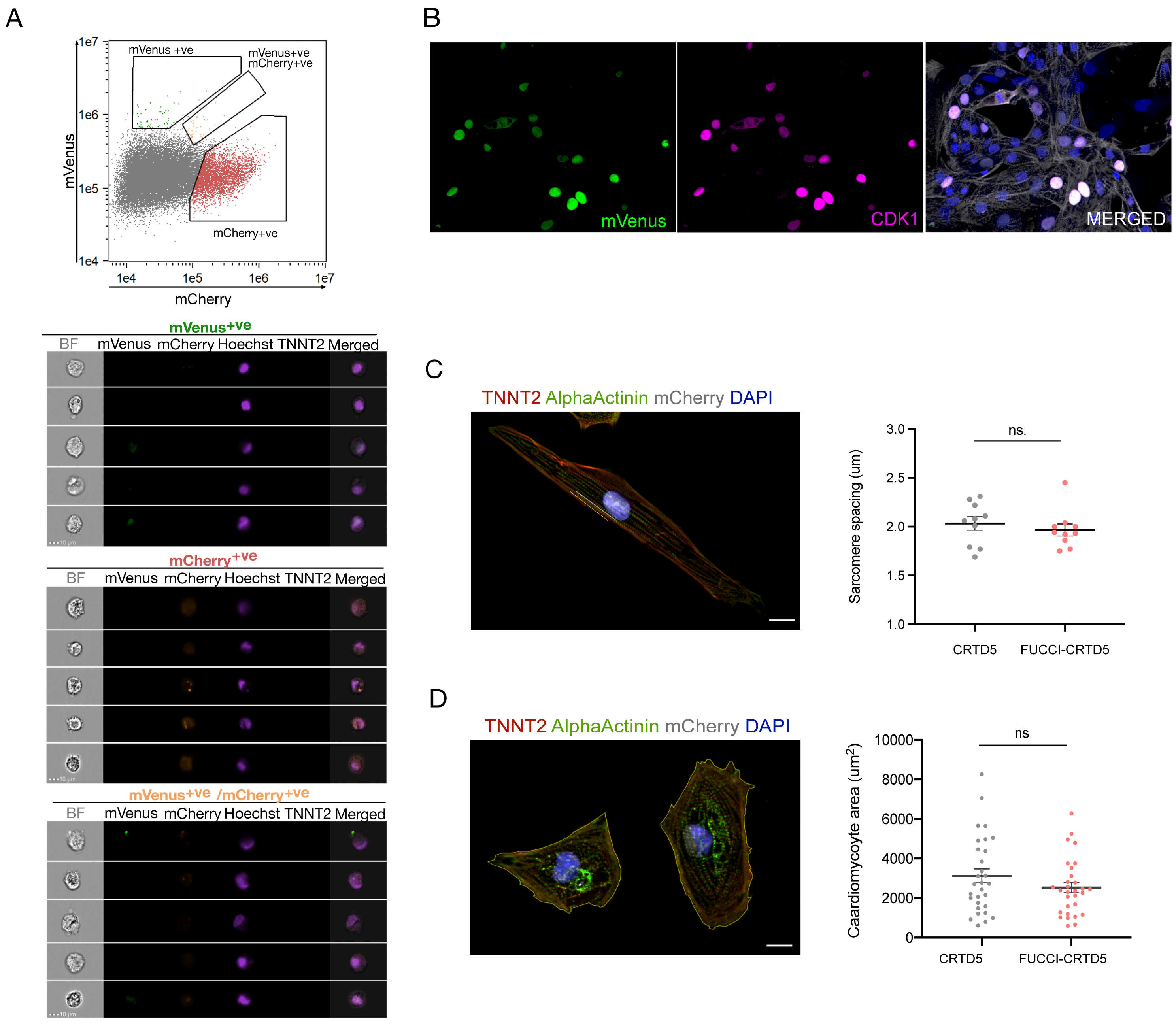


SUPPLEMENTARY FIGURE 2

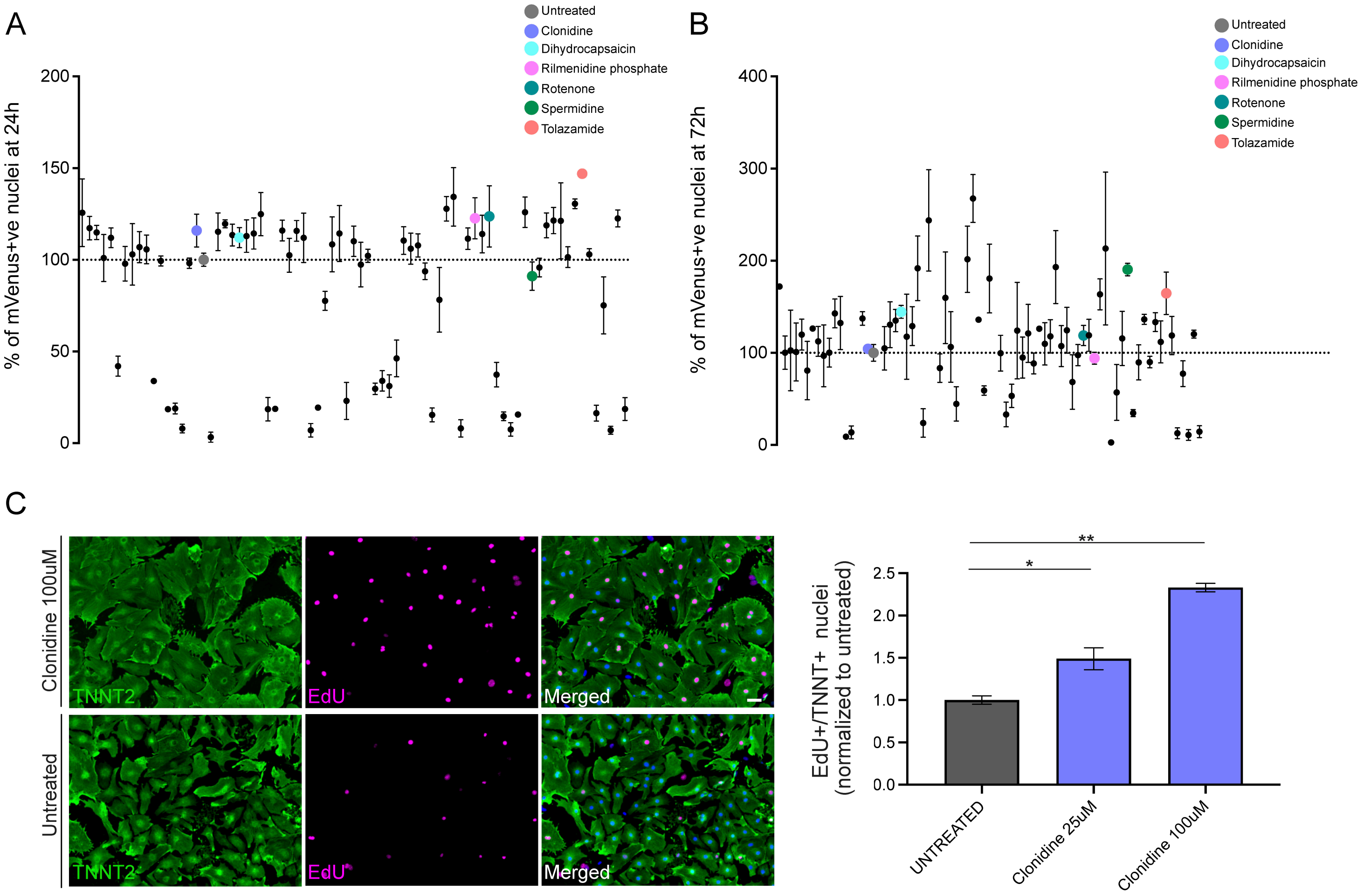


SUPPLEMENTARY FIGURE 3

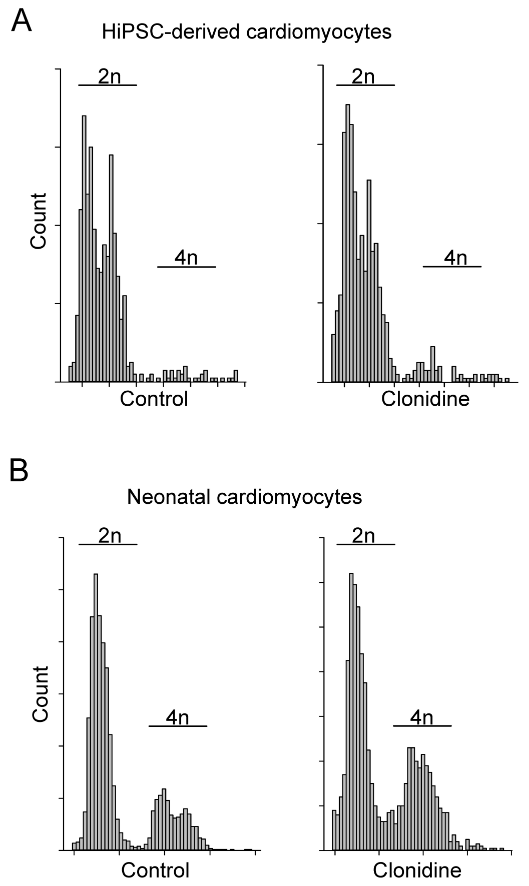


SUPPLEMENTARY FIGURE 4

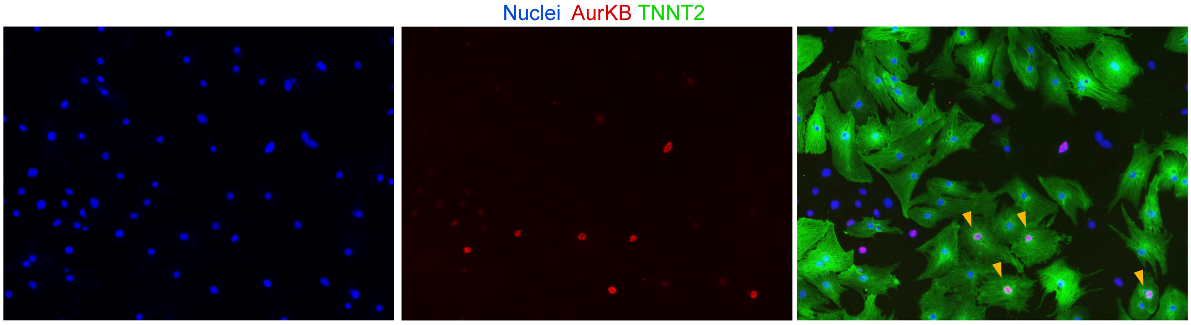


SUPPLEMENTARY FIGURE 5
