## Supplemental Table 1 for "FUCCI-based live imaging platform reveals cell cycle dynamics and identifies pro-proliferative compounds in human iPSC-derived cardiomyocytes"

| IUPAC Name | Name | Compound Purpose | Target/Mode of Action |
|---|---|---|---|
| N-(2,6-dichlorophenyl)-4,5-dihydro-1H-imidazol-2-amine hydrochloride | Clonidine. hydrochloride | Autophagy inducer | I1R agonist, ↓cAMP |
| N-(dicyclopropylmethyl)-4,5-dihydro-1,3-oxazol-2-amine;phosphoric acid | Rilmenidine phosphate | Autophagy inducer | I1R agonist, ↓cAMP |
| (4-aminobutyl)(3-aminopropyl)amine | Spermidine | Autophagy inducer | ↓HATs |
| N-[(4-hydroxy-3-methoxyphenyl)methyl]-8-methylnonanamide | Dihydrocapsaicin | Autophagy inducer | ROS accumulation |
| 1-(azepan-1-yl)-3-(4-methylbenzenesulfonyl)urea | Tolazamide | Autophagy inhibitor | ATP-K+ channel antagonist |
| (1S,6R,13S)-16,17-dimethoxy-6-(prop-1-en-2-yl)-2,7,20-trioxapentacyclo[$11.8.0.0^{3,11}.0^{4,8}.0^{14,19}$]henicosa-3(11),4(8),9,14(19),15,17-hexaen-12-one | Rotenone | Autophagy inducer | mETC-complex1-inhib |

Compound information as presented in the SCREEN-WELL® Autophagy library by ENZO life sciences
